# Novel Mouse Model of Alternating Hemiplegia of Childhood Exhibits Prominent Motor and Seizure Phenotypes

**DOI:** 10.1101/2024.08.18.608455

**Authors:** Nicole A. Hawkins, Jean-Marc DeKeyser, Jennifer A. Kearney, Alfred L. George

## Abstract

Pathogenic variants in *ATP1A3* encoding the neuronal Na/K-ATPase cause a spectrum of neurodevelopmental disorders including alternating hemiplegia of childhood (AHC). Three recurrent *ATP1A3* variants are associated with approximately half of known AHC cases and mouse models of two of these variants (p.D801N, p.E815K) replicated key features of the human disorder, which include paroxysmal hemiplegia, dystonia and seizures. Epilepsy occurs in 40-50% of individuals affected with AHC, but detailed investigations of seizure phenotypes were limited in the previously reported mouse models. Using gene editing, we generated a novel AHC mouse expressing the third most recurrent *ATP1A3* variant (p.G947R) to model neurological phenotypes of the disorder. Heterozygous *Atp1a3*-G947R (*Atp1a3*^G947R^) mice on a pure C57BL/6J background were born at a significantly lower frequency than wildtype (WT) littermates, but *in vitro* fertilization or outcrossing to a different strain (C3HeB/FeJ) generated offspring at near-Mendelian genotype ratios, suggesting a defect in reproductive fitness rather than embryonic lethality. Heterozygous mutant mice were noticeably smaller and exhibited premature lethality, hyperactivity, anxiety-like behaviors, severe motor dysfunction including low grip strength, impaired coordination with abnormal gait and balance, and cooling-induced hemiplegia and dystonia. We also observed a prominent seizure phenotype with lower thresholds to chemically (flurothyl, kainic acid) and electrically induced seizures, post-handling seizures, sudden death following seizures, and abnormal EEG activity. Together, our findings support face validity of a novel AHC mouse model with quantifiable traits including co-morbid epilepsy that will be useful as an *in vivo* platform for investigating pathophysiology and testing new therapeutic strategies for this rare neurodevelopmental disorder.

## INTRODUCTION

Alternating hemiplegia of childhood (AHC) is a rare neurodevelopmental disorder most often associated with pathogenic missense variants in *ATP1A3* encoding the neuronal sodium-potassium adenosine triphosphatase (Na/K-ATPase) (1, 2). Onset of AHC occurs typically during infancy with bouts of abnormal eye movements, paroxysmal hemiplegia, hemiparesis or quadriparesis lasting minutes to days, and dystonia (3-5). Developmental delay and intellectual disability are common, as are other nonparoxysmal features such as hypotonia and abnormal gait. Importantly, while AHC is named for the prominent movement disorder, approximately 50% of individuals with AHC have epilepsy starting either before or after onset of motor symptoms (6). Moreover, seizures can be the presenting manifestation of AHC. This disorder can be progressive and disabling, and there is no cure or universally effective treatment. A variety of other conditions have been reported in association with pathogenic *ATP1A3* variants suggesting that *ATP1A3*-related disorders encompasses a continuum of neurodevelopmental and neurological disease with overlapping clinical features (7-9), which may suggest shared pathogenic mechanisms.

There are three recurrent missense *ATP1A3* variants associated with AHC that account for approximately half of all genotype-known cases. The two most recurrent variants discovered in more than one third of genotype-positive individuals are p.D801N and p.E815K, with a third variant, p.G947R, observed in approximately 10% of cases. Most evidence supports a molecular loss-of-function (1, 10, 11) but AHC likely does not arise from pure haploinsufficiency given the absence of truncating variants associated with this disease (4).

Recapitulating phenotypes in an animal model is valuable for investigating disease pathophysiology and establishing *in vivo* platforms for testing therapeutic strategies. Various mouse models with missense mutations in murine *Atp1a3* have been described (12). The first reported *Atp1a3* missense mutant mouse was discovered during an *in vivo* chemical mutagenesis screen for dominant visible phenotypes (13). One heterozygous mutant, designated as *Myshkin* (*Myk*), exhibited heritable small body size and recurrent convulsive seizures attributed to a dysfunctional missense *Atp1a3* mutation (p.I810N). Motor dysfunction was not reported initially but was later recognized following discovery of human *ATP1A3* mutations associated with AHC (14). The specific variant responsible for the *Myk* phenotype is rarely observed in association with human AHC (4).

The two most recurrent pathogenic variants associated with AHC (p.D801N, p.E815K) were modeled in mice (15, 16). Both mutations were associated with inducible hemiplegia and dystonia, abnormal motor performance, and spontaneous or inducible seizures in heterozygous mutant mice. Each of these mouse models exhibited high mortality rates following seizures, and electrocortical recordings demonstrated abnormal activity, but quantitative analyses were limited. *ATP1A3* p.G947R, the third most recurrent variant associated with AHC, has never been modeled in mice.

In this study, we generated a novel mouse model of AHC (*Atp1a3*^G947R^) heterozygous for the recurrent p.G947R variant and performed extensive motor and seizure phenotyping. Heterozygous *Atp1a3*^G947R^ mice were smaller, had lower long-term survival, impaired motor function, greater susceptibility to seizure induction, and handling-induced and spontaneous behavioral and electrographic seizures. The availability of this mouse model provides new opportunities to investigate the pathophysiology of AHC and to establish genotype-phenotype relationships among the most recurrent *ATP1A3* pathogenic variants using *in vivo* models.

## METHODS

### Generation of an Atp1a3^G947R^ mouse model

*Atp1a3*^G947R^ (*Atp1a3*^R/+^) mice were generated using CRISPR/Cas9 editing. Guide RNA (5’-GAATAAGATCTTGATCTT-3’) and single stranded DNA donor repair oligonucleotide (Supplemental Table S1) were microinjected into C57BL/6J embryos by the Northwestern University Transgenic and Targeted Mutagenesis Laboratory (TTML). The specific introduced nucleotide variant was c.2839G>C (NM_001290469) with codon change from GGC to CGC. Founders were screened using primers outside the repair oligonucleotide homology region. Amplicons were TOPO-cloned and Sanger sequenced (primer sequences provided in Supplemental Table S1). Male mosaic founders were bred to wild-type (WT) female C57BL/6J mice (#000664, Jackson Laboratories, Bar Harbor, ME, USA) to generate N1 offspring, which were Sanger sequenced to confirm editing of the intended mutation. Predicted off-target edits with 3 or fewer mismatches with the guide RNA involving coding sequences or intronic sequences near splice sites in genes including *Atp1a3* and the related Na/K-ATPase genes *Atp1a1, Atp1a2* and *Atp1a4* were screened. Heterozygous N1 generation males with the confirmed mutation were bred with C57BL/6J females to establish the line. It was discovered during breeding of early generations that offspring genotypes were not consistent with Mendelian inheritance. Therefore, sperm was cryopreserved from sequence-verified N3-N5 generation *Atp1a3*^R/+^ males. Male and female *Atp1a3*^R/+^ and WT littermates were generated for experiments by *in vitro* fertilization (IVF). Genotypes of IVF-generated offspring were more consistent with Mendelian inheritance. To generate F1.*Atp1a3*^*R/+*^ offspring, B6.*Atp1a3*^*R/+*^ males were crossed with C3HeB/FeJ females (#000658, Jackson Laboratories). Mice were maintained in a Specific Pathogen Free (SPF) barrier facility with a 14-h light/10-h dark cycle and access to food and water *ad libitum*.

All animal care and experimental procedures were approved by the Northwestern University Animal Care and Use Committees in accordance with the National Institutes of Health Guide for the Care and Use of Laboratory Animals. The principles outlined in the ARRIVE (Animal Research: Reporting of *in vivo* Experiments) guideline were considered when planning experiments (17).

### Genotyping

Mice were genotyped by PCR using genomic DNA isolated from tail biopsies and a restriction fragment length polymorphism (RFLP) assay. Genomic DNA was amplified using RFLP genotyping primers (Supplemental Table S1), followed by restriction enzyme digest with Aci1 (#0551, New England Biolabs, Ipswich, MA, USA) for ≥ 1.5 hours at 37°C. Digestion results in 358 and 121 bp bands for WT and 358, 242, 121 and 116 bp bands for *Atp1a3*^R/+^.

### Cohorts and statistical analyses

Equal sized groups of male and female WT and *Atp1a3*^R/+^ littermates of similar age were used for all experiments (Supplemental Table S2). Groups were collapsed by sex for analysis unless specified. Data were analyzed in real-time or offline by a reviewer blinded to genotype, although blinding was confounded for some experiments due to overt phenotypes. Normality was assessed by the D’Agostino & Pearson test for all experiments. Normality and statistical comparisons were determined using GraphPad Prism v10.2.1. All statistical tests, p-value thresholds and *post hoc* analyses are presented in Supplemental Table S3. Initial comparisons were made between the sexes within groups. There were no significant differences between sexes on reported measures, except for body weight and length. Therefore, for all other measures, groups were collapsed across sex for analysis.

### Expression analysis

Mouse forebrain tissues were isolated and dissected at the midline to create equal halves for transcript and protein analysis. Total RNA was extracted using TRIZol (Invitrogen, Waltham, MA, USA) according to the manufacturer’s instructions. First strand cDNA was synthesized using 4 μg of total RNA using oligo(dT) primer and Superscript IV reverse transcriptase (RT; Invitrogen) according to the manufacturer’s instructions. First strand cDNA was diluted 1:1000 and 5 μL was used as template in droplet digital PCR (ddPCR) using ddPCR Supermix for Probes (no dUTP; Bio-Rad, Des Plaines, IL, USA) and TaqMan Gene Expression Assay probes (Applied Biosystems, Waltham, MA) for mouse *Atp1a3* (FAM-MGB-Mm00523430_m1) and *Gapdh* (normalization control; VIC-MGB-Mm99999915_g1). Reactions were partitioned using a QX200 droplet generator (Bio-Rad) and then amplified using PCR conditions: 95°C for 10 min, 44 cycles of 95°C for 30 s and 60°C for 1 min (ramp rate of 2°C/s) with a final inactivation step of 98°C for 5 min. Following amplification, droplets were analyzed with a QX200 droplet reader and QuantaSoft vl.6.6 software (Bio-Rad). Relative transcript levels were expressed as a ratio of *Atp1a3* to *Gapdh* normalized to WT and included 9–11 biological replicates per genotype. Both assays lacked detectable signal in genomic, no-RT and no-template control reactions.

Membrane proteins were isolated using P3 membrane protein fractionation. Samples (30 μg) were size-separated on 7.5% SDS-PAGE gels and transferred to nitrocellulose. Blots were probed with anti-ATP1A3 (1 μg/mL; XVIF9-G10, Abcam, Cambridge, UK) and anti-mortalin/GRP75 (1:1000; 75-127, NeuroMab, Davis, CA, USA) antibodies. Alexa-conjugated fluorescent anti-mouse 790 secondary antibody was used to detect signal using an Odyssey imaging system (LI-COR, Lincoln, NE, USA). Relative protein levels were determined by densitometry using Image Studio (LI-COR) and expressed as a ratio of Atp1a3 to GRP75 normalized to WT. We assayed 8-11 biological replicates per genotype.

### Body size and weight

Mice were weighed twice between 8-16 weeks of age with 43-51 mice of each sex per genotype. Body length from end of tail to tip of nose was measured once at 8-16 weeks of age, with 20-26 of each sex per genotype.

### Survival Analysis

Mice were weaned and survival was monitored to approximately 5 months of age. Survival was compared between genotypes by Kaplan Meier analysis (161-178 mice per genotype).

### Open field

Mice were tested separately by sex with at least a 1-hour delay between sexes with 15-18 of each sex per genotype. Prior to testing, mice were acclimated to the testing suite with white noise for 1 hour. Mice were placed in the center of an open field arena (46 x 46 cm) and video monitored for 30 minutes. Limelight software (Actimetrics, Wilmette, IL, USA) was used to acquire trial data. Ethovision XT 15 (Noldus, Leesburg, VA, USA) was used to analyze distance traveled and % time spent in the arena center.

### Response to tail suspension

Mice were tested separately by sex with at least a 1-hour delay between groups, with 14-19 of each sex per genotype. Prior to testing, mice were acclimated to the testing suite with white noise for 1 hour. Mice were suspended by their tails for 1 min, then were scored using a lab-developed rating scale based on observable behaviors that included movement abilities (Score: 1 - immobile hang, 2 - tremor, 3 -spastic/erratic), limb dysfunction (Score: 1 - none, 2 - hindlimb/forelimb clasp and/or paddle, 3 - unilateral movement) and climbing behaviors (Score: 1 - able to curl/climb up tail to right self, 2 - trying to curl up and right self unsuccessfully, 3 - unable to curl up). Mice were tested over two consecutive days and cumulative scores (range 3 - 9) were averaged with date from days.

### Forelimb adhesive removal assay

Mice were tested separately by sex with at least a 1-hour delay between groups, with 14-19 of each sex per genotype. Prior to testing, mice were acclimated to the testing suite with white noise for 1 hour. An adhesive sticker (8 x 12 mm) was placed on the head of each mouse, and the mouse was returned to an observation cage with only bedding. Latency to remove the sticker with forelimbs was recorded. Mice were tested over two consecutive days and latency was averaged with data from both days.

### Performance on elevated balance beam

Mice were tested separately by sex with at least a 1-hour delay between groups, with 13-19 of each sex per genotype. Prior to testing, mice were acclimated to the testing suite with white noise for 1 hour. Mice were placed on the beam and allowed to traverse the length. Latency to cross or fall was recorded. Mice were tested over three consecutive days and latency was averaged for all days.

### Performance on elevated wire hang

Mice were tested separately by sex with at least a 1-hour delay between groups, with 14-19 of each sex per genotype. Prior to testing, mice were acclimated to the testing suite with white noise for 1 hour. Mice were placed on their home cage wire rack which was slowly inverted and raised to a height of 23 cm. Latency to fall was recorded, with a maximum grasp and hold time allowed of 3 minutes. Mice were tested over two consecutive days and latency was averaged with data from both days.

### Accelerating rotarod assay

Mice were tested on an accelerating rotarod (Panlab, Harvard Apparatus, Barcelona, Spain). Up to 5 mice were placed on the rotating rod, separated by dividers. Rotation was accelerated from 4 to 40 RPM over a 5 min session. Mice were given three trials per day for 3 consecutive days with an inter-trial interval of 15 min. Latency to fall from the rotating rod was automatically recorded. A daily average was calculated for each mouse from the three trials. Average latencies on the final day were compared between genotypes using an unpaired t-test.

### Gait analysis

Fore and hind paws were dipped in a non-toxic fingerpaint and mice were placed on a strip of paper between two guide walls leading to a dark chamber. Distance between successive prints (stride) and distance between each pair of prints (base) were measured separately for the forepaws (orange paint) and hindpaws (blue paint), as well as overlap of fore and hind paw prints on the left and right sides. Parameters were compared between groups using two-way ANOVA with Sidak’s post-hoc test.

### Hypothermia-induced hemiplegia and dystonia

Mice were monitored continually for core body temperature using a rectal thermal probe and TCAT-2DF temperature controller (Physitemp, Clifton, NJ, USA). Body temperature was slowly elevated from baseline to 42.0°C and then mice were immediately transferred to a shallow ice water bath (24 x 19 cm). The ice water level was maintained at ∼ 3 cm in depth and <2°C, to allow for rapid cooling, yet shallow enough mice could touch without forced swimming. Mice were maintained in the ice bath until a core body temperature of 30°C was reached or until dystonia-induced submersion occurred. Once removed from ice bath, mice were allowed to recover on a 42.0°C water recirculating heating pad for at least 5 minutes. If the mouse appeared abnormal (tremors, dystonia, immobility, circling, head bobbing) after the 5 min period, the mouse was video monitored until normal behaviors resumed (cage exploration, grooming). Mice were scored on a lab-developed movement disorder rating scale based on observable behaviors during the heating, cooling and recovery phases of the assay with 20 mice per genotype (Table 1). Scores were totaled (0-16) per mouse, and time to cool and recover were recorded.

**Table 1.** Movement disorder rating scale.

| **Heating Phase** |  |
| --- | --- |
| 0 | Jumping and/or escape behavior |
| 1 | Escape behaviors that decline with increased body temperature, >41°C little to no movement |
| 2 | Head bob and/or tremors |
| **Cooling Phase** |  |
| *Ice Bath Entry* |  |
| 0 | Stationary |
| 1 | Wild, erratic or uncoordinated |
| 2 | Dystonia and/or paresis |
| *Mobility* |  |
| 0 | Escape behaviors; Jumping walls, circling perimeter |
| 1 | Exploration with periods of slow mobility or immobility |
| 2 | Dystonia and/or paresis |
| *Behavior* |  |
| 0 | Unremarkable |
| 1 | Vocalizations |
| 2 | Head bob and/or tremors |
| *Duration* |  |
| 0 | Reach 30°C |
| 1 | Water submersion, removed from ice bath, recovered |
| 2 | Water submersion, removed from ice bath, dead |
| **Recovery Phase** |  |
| *Mobility & Appearance* |  |
| 0 | Mobile, unremarkable |
| 1 | Immobile or slow mobility, will move when touched |
| 2 | Dystonia; loss of posture, no control over limbs, hindlimb and/or forelimbs splayed out, neck and/or core twisted, no motor control |
| *Seizure-like* |  |
| 0 | None |
| 1 | Tail Straub |
| 2 | Head bob and/or tremors |
| *Righting behaviors* |  |
| 0 | Immediate |
| 1 | Slow/delayed |
| 2 | Unable; stays on back |

### Post-handling seizure monitoring

Mice ranging from P37-40 were video-monitored from above for one hour to capture post-handling seizures. Three to four mice of mixed genotype and sex were placed in monitoring cages with *ad libitum* access to food and water. Videos were scored offline by a blinded reviewer. Differences in seizure frequency were determined by unpaired t-test and % of mice with seizures was compared using Fisher’s exact test with 13 mice per genotype.

### Flurothyl induced seizures

Mice were exposed to the chemoconvulsant flurothyl (Bis[2,2,2-trifluoroethyl] ether, Sigma-Aldrich, St. Louis, MO, USA). Mice were placed in a Plexiglas chamber (2.2 L) and flurothyl was introduced using a syringe pump (20 uL/min) and allowed to volatilize. Latencies to the first myoclonic jerk (MJ) and onset of a generalized tonic-clonic seizure (GTCS) were recorded, with 19-20 mice per genotype. Statistical differences in latencies were determined by Mann-Whitney test.

### Electroconvulsive shock (ECS)

Mice (9-10 per genotype) were stimulated with corneal electrodes following application of 0.5% tetracaine to both eyes. Both sexes received a subthreshold shock of 60 Hz/0.5 pulse width/0.2 sec/38 mA. Seizures were scored for absence or presence of a tonic hindlimb extension (HLE) seizure within 5 seconds. Statistical differences in seizure susceptibility were determined by Fisher’s exact test.

### Kainic acid induced seizures

Kainic acid (KA, Tocris, Bristol, UK) was administered intraperitoneally at 25 mg/kg to 16-22 mice per genotype which were then video recorded for 2 hours. Videos were scored offline with Ethovision XT 15 using a modified Racine scale (18) (1: behavioral arrest; 2: Straub tail, automatisms - facial and/or repetitive scratching, circling, forelimb clonus without falling; 3: forelimb clonus with rearing and/or falling; 4: GTCS with wild running and/or jumping; 5: death). Latency to each stage was recorded. Data were binned into 5 min segments and the highest stage reached was recorded. Statistical differences for KA stage latencies were determined by Two-Way ANOVA with Sidak’s *post hoc* tests and survival was determined by Mantel-Cox.

### Video-EEG monitoring

Mice were implanted with prefabricated 2 EEG/1 EMG channel headmounts (8201; Pinnacle Technology, Lawrence, KA, USA). Mice were anesthetized with ketamine/xylazine and four stainless steel screws that served as cortical surface electrodes and the headmount were affixed to the skull with glass ionomer cement (GC FujiCEM, GC Dental, Alsip, IL, USA). Anterior screw electrodes were 0.5–1 mm anterior to bregma and 1 mm lateral from the midline. Posterior screws were 4.5–5 mm posterior to bregma and 1 mm lateral from the midline. EEG1 represents recordings from right posterior to left posterior (interelectrode distance ≈2 mm) and EEG2 represents recordings from right anterior to left posterior (interelectrode distance ≈5 mm). The left anterior screw served as the ground connection. Electromyogram (EMG) leads were inserted into the trapezius muscle and secured with sutures. Following at least 48 hours of recovery, tethered EEG/EMG and video data (Sirenia acquisition software, Pinnacle Technology) were continuously collected for 7 days. EEG data was acquired at a sampling rate of 400 Hz. On average, ∼99 hours of EEG data were analyzed. Video-EEG records were manually reviewed for electrographic seizures (≥2x baseline; ≥10 sec; evolution in frequency and amplitude) and interictal epileptiform discharges, including spike trains and short bursts of high frequency, high amplitude activity. Spike trains were defined as having regular, rhythmic sharp spike and slow wave components, less than 3 Hz, with ≥1 spike per second continuing for ≥20 seconds. Short bursts were defined as high frequency, high amplitude activity occurring in 3-12 sec long segments. The root mean square (RMS) was calculated using LabChart v8.1.19 (Adinstruments, Sydney, Australia) for the entire 7 days and was considered baseline average for each file.

To quantify baseline EEG activity and generate spectral arrays, we used a fast Fourier transform (FFT) size of 256, 93.75% overlap window and Hann-Cosine-Bell fit, which generated approximately 22 frequency data points between 0-32 Hz (LabChart). After visual inspection, 5 min epochs of spike-, burst- and artifact-free EEG traces were selected for spectral analysis and total power data. Power spectral density (PSD) values were generated separately for epochs during periods of wakefulness and apparent sleep(absent movement and not EMG signal).

## RESULTS

### Generation of heterozygous Atp1a3-G947R mice

We generated the *Atp1a3*-G947R (*Atp1a3*^G947R^ or *Atp1a3*^R/+^) mouse model of AHC on the C57BL/6J strain using CRISPR/Cas9 gene editing. The intended mutation was verified by Sanger sequencing of genomic PCR products (Fig. 1A). The mutation was not associated with differences in brain *Atp1a3* transcript or protein levels between *Atp1a3*^R/+^ and wildtype (WT) mice (Fig. 1B,C; Supplemental Fig. S1). Screening for predicted off-target edits including *Atp1a1, Atp1a2*, and *Atp1a4*, was negative.

**Fig. 1.**
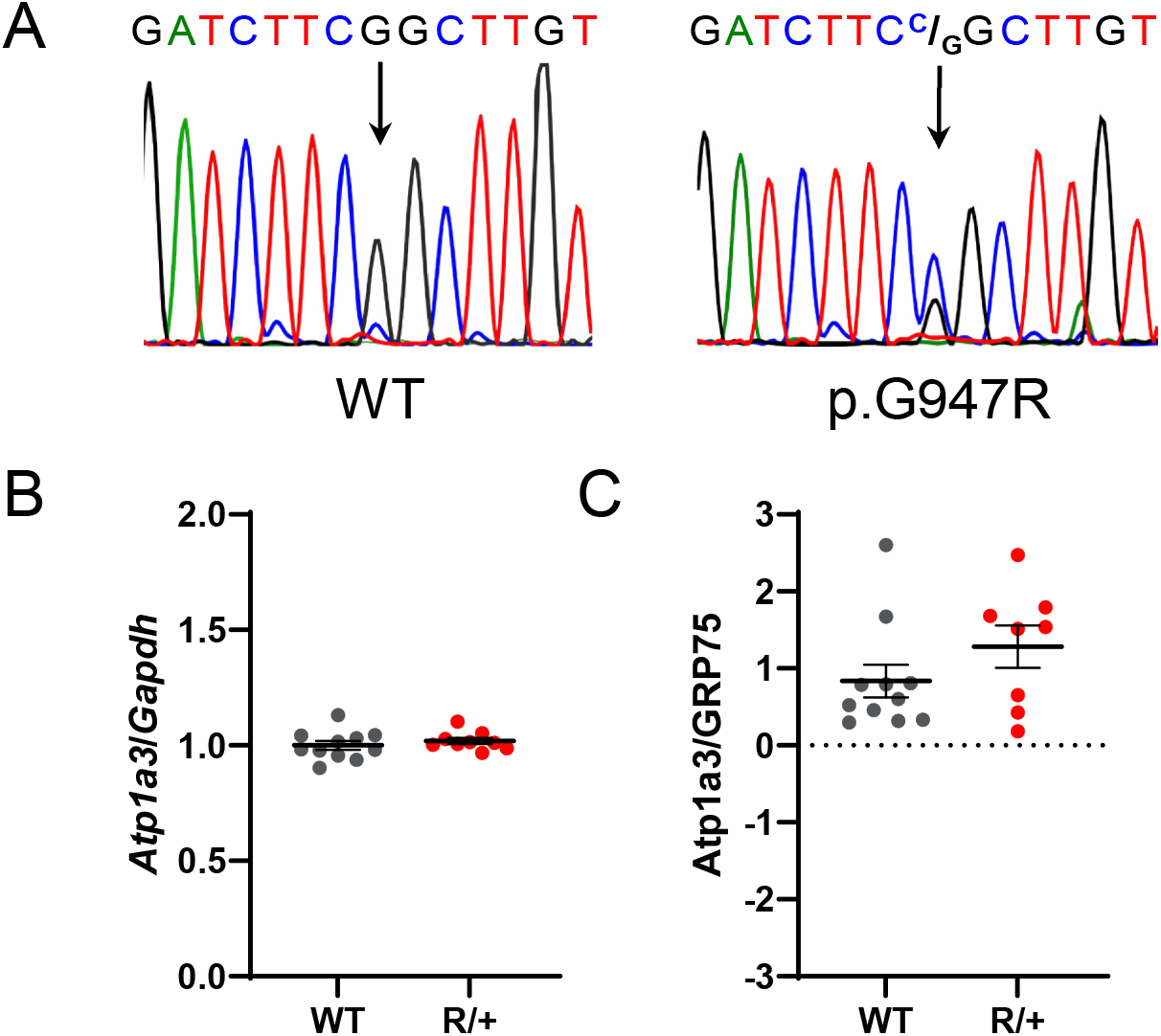
Generation and validation of the *Atp1a3*^G947R^ mouse model. **A**. Genomic amplicon sequence electropherograms comparing WT and *Atp1a3*^G947R^ (R/+) mice. **B**. *Atp1a3* mRNA transcript expression in brain normalized to *Gapdh* was not different between WT and R/+ mice (p=0.43). **C**. *Atp1a3* protein expression in brain normalized to GRP75 was not different between WT and R/+ mice (p=0.31). In panels B and C, symbols are data from individual mice, horizontal lines represent mean values and error bars indicate SEM, n = 8-11 mice per genotype.

The distribution of offspring genotypes resulting from conventional breeding of *Atp1a3*^R/+^ males with C57BL/6J females differed from expected ratios with fewer mutant mice born (Table 2). Only 5% of males and 13% of females were heterozygous for *Atp1a3*-G947R, skewing the expected WT proportion from the expected 25% per sex to 39% of males and 43% of females (p<0.0001). This breeding strategy produced typical C57BL/6J litter sizes (5.1 ± 0.6, range: 1-16, n=165 mice; 33 litters). When C57BL/6J *Atp1a3*^R/+^ males were crossed with C3HeB/FeJ females to generate F1 offspring, genotypes were more consistent with Mendelian inheritance, suggesting a potential strain-specific genetic modifier for the variant allele rather than an embryonic lethal phenotype (Table 2). To combat the low transmission rate of the variant on C57BL/6J, we used *in vitro* fertilization (IVF) to produce mutant and WT littermates for experiments. While IVF greatly improved the number of *Atp1a3*^R/+^ offspring, the observed genotype proportions still deviated slightly from the expected Mendelian ratios (p<0.04).

**Table 2.**
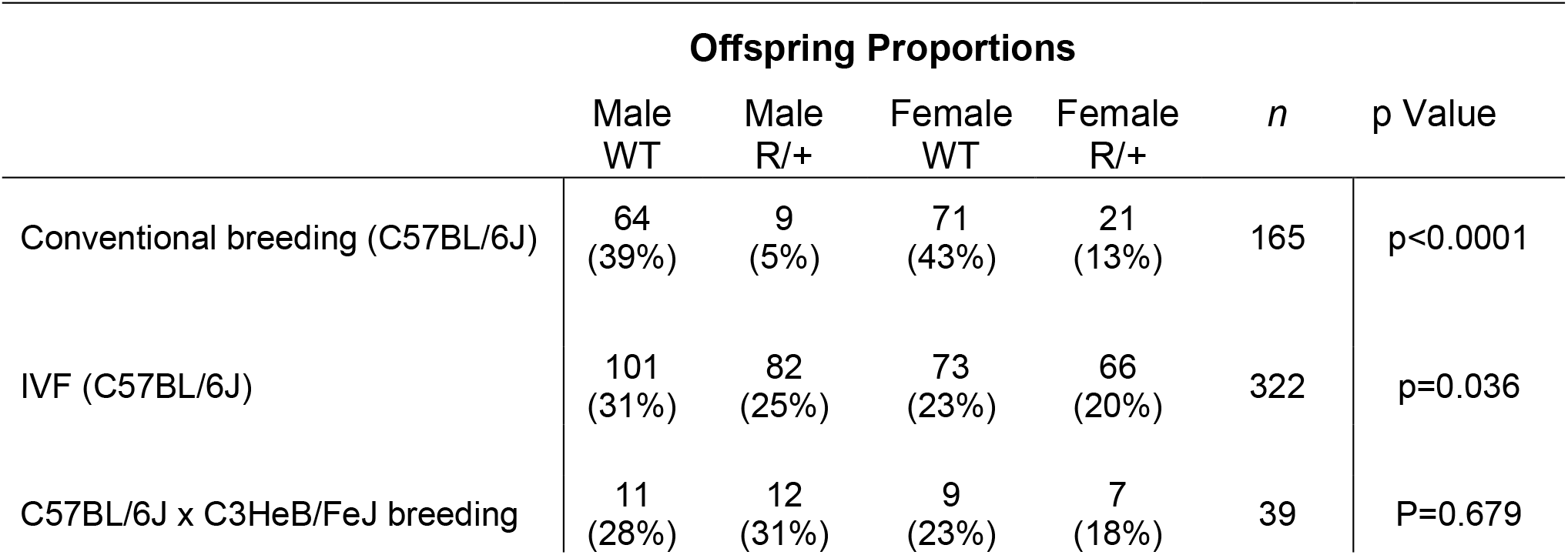
Sex and genotype ratios of offspring.

### Atp1a3-G947R impacts physical characteristics and survival

*Atp1a3*^R/+^ males and females weighed 10-20% less than their WT littermates (Fig. 2A), with an average of 23.0 ± 0.4 g and 19.1 ± 0.2 g respectively, compared to the WT averages of 28.4 ± 0.4 g and 21.9 ± 0.3 g (p<0.0001). Furthermore, *Atp1a3*^R/+^ males and females exhibited shorter body length compared to WT littermates (Fig. 2B). WT males and females had average lengths of 17.3 ± 0.1 cm and 16.5 ± 0.1 cm, respectively, while *Atp1a3*^R/+^ males and females had average lengths of 16.1 ± 0.1 cm and 15.6 ± 0.1 cm, respectively (p<0.0001). Additionally, *Atp1a3*^R/+^mice exhibited premature lethality with sporadic deaths after the first month of life and more frequent lethality between 3 and 5 months of age, resulting in ∼62% survival at 5 months of age (p<0.0001; Fig. 2C). Survival was not biased by sex nor were any of the neurological phenotypes ascertained.

**Fig. 2.**
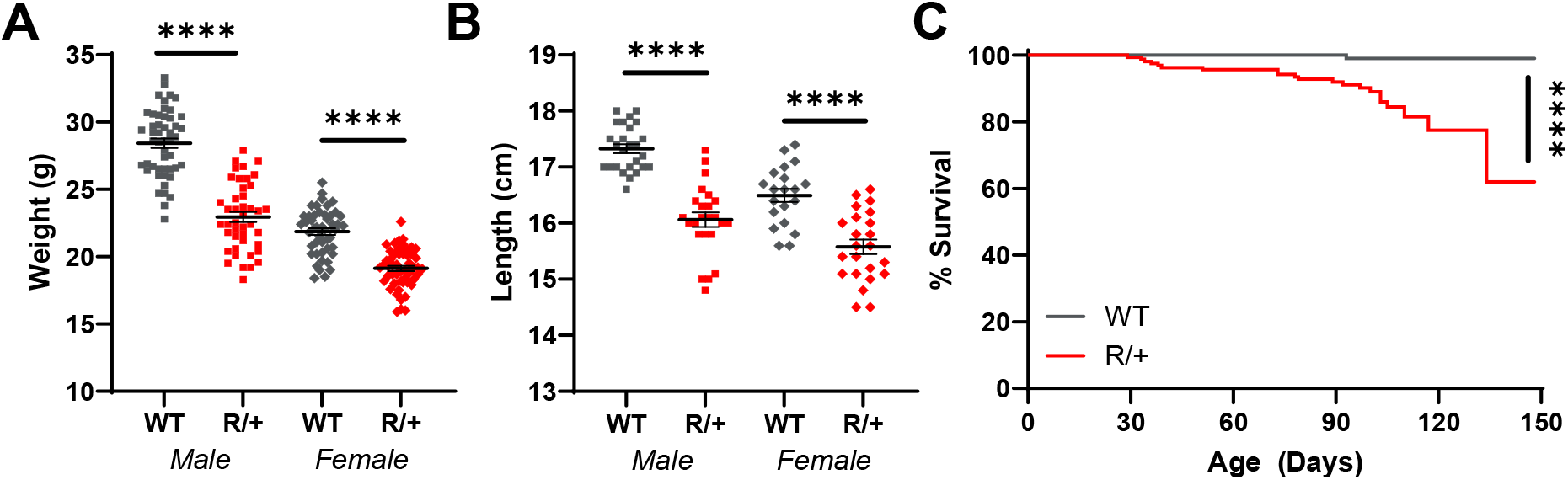
Physical characteristics and survival of heterozygous *Atp1a3*^G947R^ mice. **A**. R/+ mice weighed less than their sex-matched WT littermates. WT males and females have average weights of 28.4 ± 0.35 g and 21.9 ± 0.25 g, while R/+ males and females have average weights of 23.0 ± 0.4 g and 19.1 ± 0.2 g (****p<0.0001, 43-51 mice of each sex per genotype). **B**. R/+ mice are smaller than their sex-matched WT littermates. WT males and females have average lengths of 17.3 ± 0.1 cm and 16.5 ± 0.1 cm, while R/+ males and females have average lengths of 16.1 ± 0.1 cm and 15.6 ± 0.1 cm (****p<0.0001, 20-26 mice of each sex per genotype). **C**. R/+ mice exhibit premature lethality. Kaplan Meier plot comparing WT and R/+ survival. R/+ mice begin to sporadically die after the first month of life with increasing mortality during 3 to 5 months of life (****p<0.0001, 161-178 mice per genotype). In panels A and B, symbols represent individual mice, horizontal lines represent mean values and error bars indicate SEM.

### Atp1a3^G947R^ mice exhibit neurological and motor phenotypes

WT and *Atp1a3*^*R/+*^ mice underwent testing to evaluate motor function. In an open field arena, *Atp1a3*^*R/+*^ mice exhibited locomotor hyperactivity with an average traveled distance of 180.1 ± 10.8 m, while WT mice traveled an average distance of 130.3 ± 4.6 m (Fig. 3A; p<0.0001). However, *Atp1a3*^*R/+*^ spent a smaller percentage of time (10.4 ± 1.0%) in the center of the arena compared to WT mice (13.4 ± 0.9 %), suggestive of an anxiety-like phenotype rather than reduced exploration (Fig. 3B; p<0.03).

**Fig. 3.**
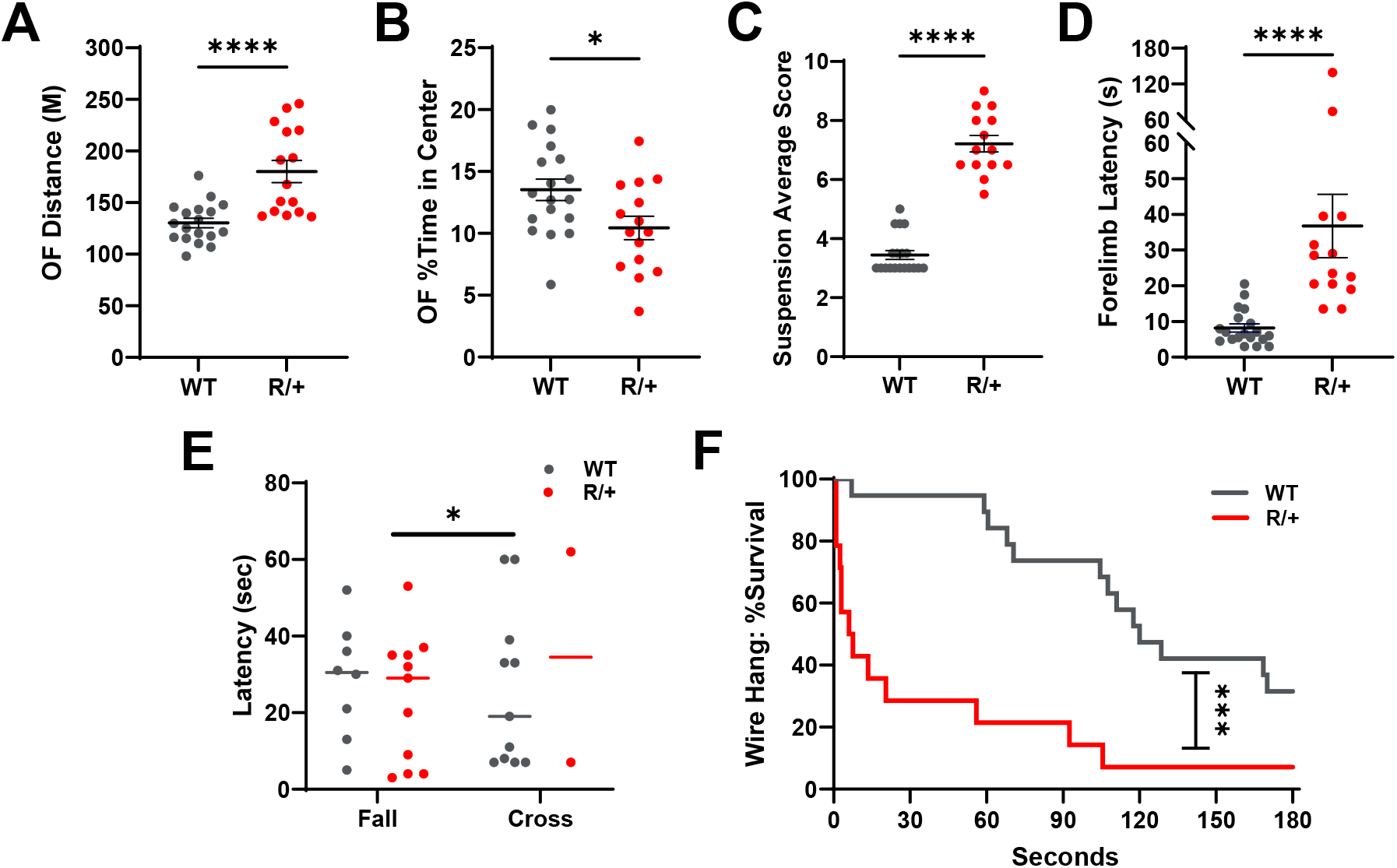
Impaired motor function of heterozygous *Atp1a3*^G947R^ mice. **A**. R/+ mice traveled farther in an open field arena compared to WT (R/+: 180.1±10.8 m; WT: 130.3±4.6 m; ****p<0.0001, 15-18 mice per genotype). **B**. R/+ mice spent less time in the center of an open field arena compared to WT (R/+: 10.4 ± 0.95%; WT: 13.4 ± 0.86%; *p<0.03, 15-18 mice per genotype). **C**. Motor behaviors during tail suspension. WT mice had an average tail suspension score of 3.45 ± 0.15, due to spending the most time in an immobile hang position or trying to right themselves. R/+ mice displayed uncoordinated and erratic trunk and limb movements, had trouble curling to right themselves and scored an average of 7.2 ± 0.28 (****p<0.0001, 14-19 mice per genotype). **D**. Forelimb motor task. WT mice exhibited an average latency of 8.2 ± 1.1 s to remove an adhesive sticker from their head using forelimbs, while R/+ mice had a longer average latency of 36.7 ± 8.9 s (****p<0.0001, 14-19 mice per genotype). **E**. Success in traversing an elevated beam. Most WT mice (11/19; 58.8%) traversed the beam with a median time of 19 s. Only 15.4% (2/13) of R/+ mice traversed the beam, with a median time of 34.5 s. The median fall latency was similar between WT (30.5 s) and R/+ (29.0 s; *p<0.03, 13-19 mice per genotype). **F**. Latency to fall from an elevated wire. WT had a median fall latency of 120 s while R/+ mice had a median fall latency of 6.75 s. ***p<0.0002. n= 14-19/genotype. In panels A-D, symbols are data from individual mice, horizontal lines represent mean values and error bars indicate SEM. In panel E, horizontal lines represent median values without error bars.

To further assess motor behaviors, mice were suspended by their tails for 1 min and scored using a cumulative rating scale based on general movement abilities, limb dysfunction, and climbing behavior. Relative to WT, *Atp1a3*^*R/+*^ mice exhibited severe motor dysfunction resulting in an average score of 7.2 ± 0.3 compared to 3.4 ± 0.2 for WT mice (Fig. 3C; p<0.0001). *Atp1a3*^*R/+*^ mice displayed frequent bouts of dystonic-like movements, which included uncoordinated and erratic movement of the trunk, forelimb and/or hindlimb clasping, and no ability to curl up and climb their tail in an effort to right themselves. Conversely, WT mice spent their time either in an immobile hanging position or successfully righting themselves by tail climbing, typical of the C57BL/6J strain (19).

To evaluate fine and skilled motor function, we tested mice using the adhesive removal task (20, 21). WT mice were able to quickly swipe a sticker off their head using their forelimbs, with an average latency of 8.2 ± 1.1 s (Fig. 3D). In contrast, *Atp1a3*^*R/+*^ mice exhibited delayed forelimb motor function, with an average removal latency of 36.7 ± 8.9 s (p<0.0001).

Gait, balance, and motor coordination were also abnormal in *Atp1a3*^*R/+*^ mice. On an elevated balance beam, *Atp1a3*^*R/+*^ mice were often observed to be stuck straddling the beam or attempting to drag across using only forelimbs. While the latency to fall was not different between genotypes, only 15.4% (2/13, p<0.03) of *Atp1a3*^*R/+*^ mice successfully traversed the elevated balance beam in a median time of 34.5 s compared to 19 s for WT mice (Fig. 3E). Grip strength was assessed by placing mice on their home cage wire rack then slowly inverting and elevating the rack. Only a single (7%, 1/14) *Atp1a3*^*R/+*^ mouse was able to maintain grip for 3 minutes as compared to 32% (6/19) of WT mice (Fig. 3F). *Atp1a3*^*R/+*^ mice exhibited a rapid median fall latency of 6.8 sec compared to 120 sec for WT (p<0.0002). We identified gait abnormalities in *Atp1a3*^*R/+*^ mice, including reduced forelimb (p<0.02) and hindlimb (p<0.03) stride distance compared to WT (Supplemental Fig. S2A,B). On an accelerating rotarod, *Atp1a3*^*R/+*^ mice had a longer latency (p<0.005) to fall on all three trials compared to WT (Supplemental Fig. S2C,D). Body size was not taken into consideration for gait or rotarod analyses, and the observed differences may be a result of smaller body size (22) or hyperactivity.

### Inducible hemiplegia and dystonic-like behaviors in Atp1a3^G947R^ mice

Recurrent episodes of hemiplegia and dystonia in individuals with AHC are often provoked with exposure to water, extreme temperatures and/or stress (23). Therefore, we tested if fluctuations in body temperature provoked hemiplegia and dystonia-like behaviors in *Atp1a3*^*R/+*^ and WT mice. First, core body temperature was raised to 42°C (heating phase), followed by rapid cooling in an ice bath to 30°C (cooling phase). Following cooling and removal from the ice bath, mice were monitored for at least 5 minutes on a 42°C heating pad to observe mobility, posture/appearance and the ability to right (recovery phase). Immobility, hemiplegia and dystonic-like behaviors in *Atp1a3*^*R/+*^ mice became apparent during the cooling and recovery phases of the assay (Fig. 4A). Each mouse received a score, based on a scale focused on observable abnormal motor behaviors during the heating, cooling and recovery phases of the assay (Table 1). WT mice had an average score of 0.9 ± 0.2, while *Atp1a3*^*R/+*^ mice had an average score of 9.6 ± 0.7 (p<0.0001; Fig. 4B). Whereas all WT mice (20/20) were cooled from 42° to 30°C without exhibiting any abnormal movement behaviors, 35% (7/20) of *Atp1a3*^*R/+*^ mice exhibited severe dystonic-like behaviors or paresis and were removed promptly from the ice bath to avoid submersion. Four *Atp1a3*^*R/+*^ mice failed to recover and died during the recovery phase. During the recovery phase, all WT mice (20/20) exhibited normal mobility, body posture/appearance and were able to right themselves from a supine position (Fig. 4C,D). By contrast, none of the *Atp1a3*^*R/+*^ mice (0/16) exhibited normal recovery (p<0.0001). Instead, *Atp1a3*^*R/+*^ mice exhibited dystonic-like postures, including core and body twisting, tail rigidity, loss of posture, lack of forelimb and/or hindlimb control, and required a median recovery time of 41.4 min before regaining a normal righting reflex (Fig. 4C,E,F). In addition, many mice exhibited hemiplegia with or without dystonia during the recovery phase.

**Fig. 4.**
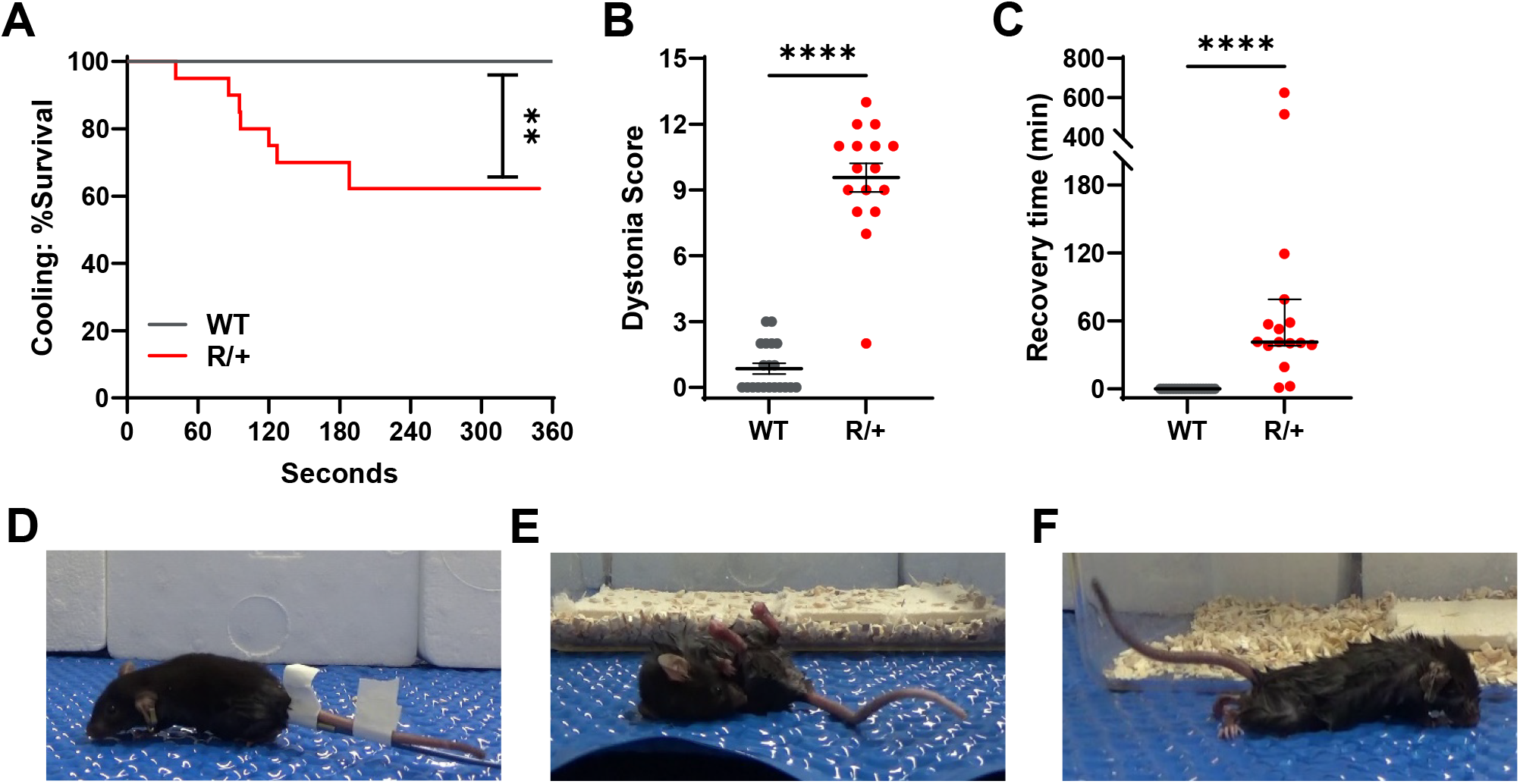
Heterozygous *Atp1a3*^G947R^ mice exhibit dystonic-like behaviors. **A**. R/+ mice became dystonic during rapid cooling. Kaplan-Meier curves for occurrence of dystonic-like behaviors upon cooling from 42° to 30°C core temperature (**p<0.004, 20 mice per genotype). **B**. Comparison of dystonia score between WT and R/+ mice (****p<0.0001, 16-20 mice per genotype). Symbols are data from individual mice, horizontal line represents mean values and error bars indicate SEM **C**. Comparison of latency to recovery from dystonic behaviors between WT and R/+ mice (****p<0.0001, 16-20 mice per genotype). Symbols are data from individual mice, horizontal lines and error bars represent median values with 95% CI. **D**. Representative image of a WT mouse, exhibiting normal appearance and mobility after cooling. **E**. Representative image of a R/+ mouse, exhibiting inability to right self, tail rigidity, hindlimb clasping, and core/body twisting after cooling. **F**. Representative image of a separate R/+ mouse, exhibiting hyperextension and splaying of hindlimbs with tail rigidity after cooling.

### Atp1a3^G947R^ mice have greater susceptibility to induced seizures

We used multiple chemical and electrical seizure-induction methods to evaluate seizure susceptibility in *Atp1a3*^*R/+*^ mice. After exposure to flurothyl, *Atp1a3*^*R/+*^ mice had an average latency to GTCS of 136.5 ± 8.5 sec, while average latency for WT mice was 181.1 ± 6.2 sec (p=0.0001; Fig. 5A). Following the onset of seizure activity, most *Atp1a3*^*R/+*^ mice displayed dystonic-like behaviors that were never observed in WT mice. Furthermore, *Atp1a3*^*R/+*^ mice transitioned more quickly between the onset of the first sign of seizure onset (myoclonic jerks) and GTCS compared to WT (p<0.0001; Fig. 5B).

**Fig. 5.**
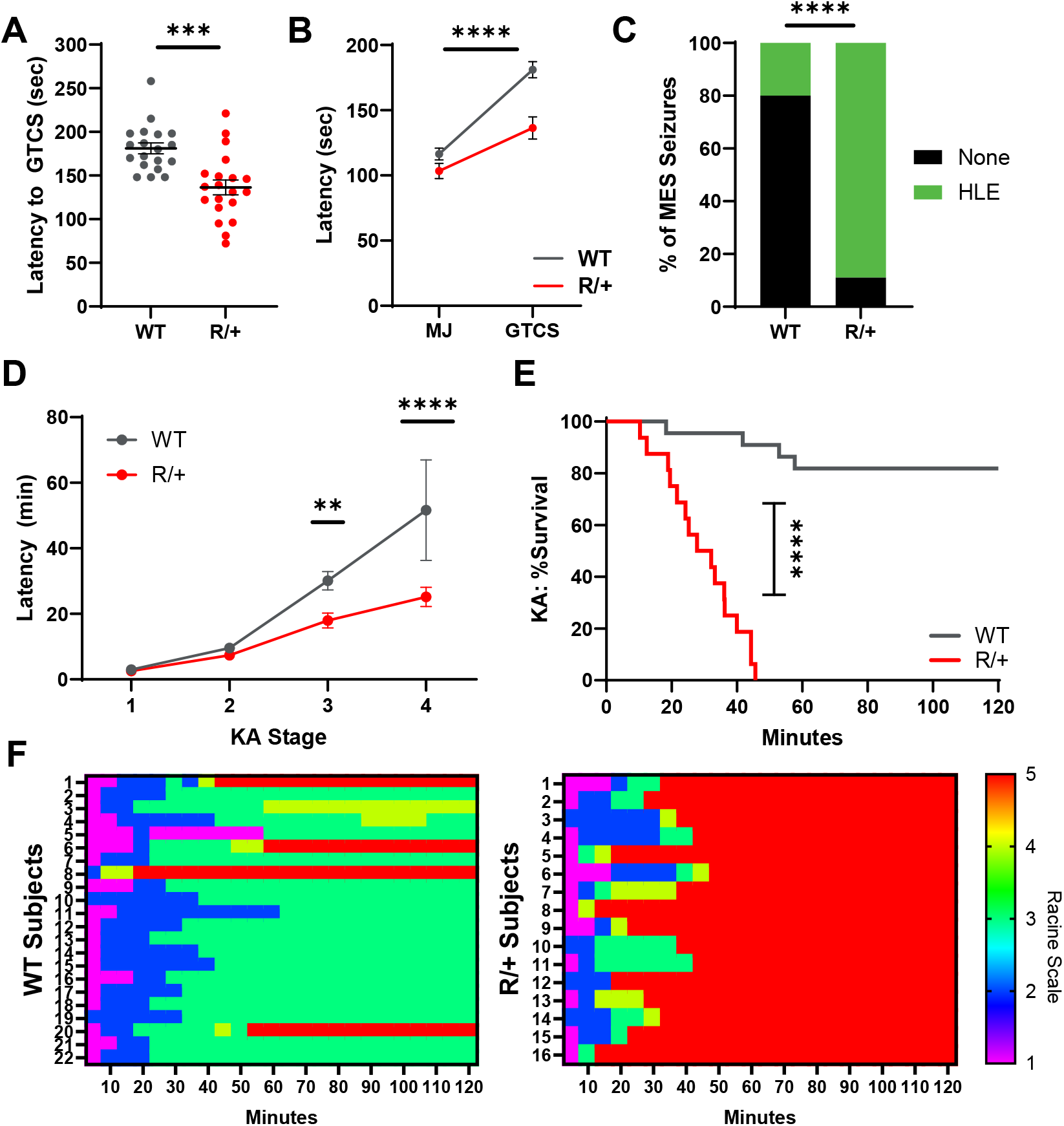
Heterozygous *Atp1a3*^G947R^ mice have lower induced seizure threshold. **A**. Comparison of latency to GTCS following flurothyl exposure between WT and R/+ mice (***p=0.0001, 19-20 mice per genotype). Symbols are data from individual mice, horizontal lines represent mean values and error bars indicate SEM. **B**. Latency to myotonic jerk (MJ) and GTCS after flurothyl exposure. There was no difference in latency to onset of MJ between WT and R/+ mice, but mutant mice transitioned more quickly to GTCS (****p<0.0001, 15-19 per genotype). **C**. Susceptibility to ECS-induced seizures. A greater proportion of R/+ mice exhibited hindlimb extension (HLE) in response to a subthreshold corneal ECS stimulation (****p<0.0001, 9-10 mice per genotype). **D**. Comparison of latency to KA induced seizures between WT and R/+ mice. Mutant mice progressed faster to Racine stages 3 and 4 (**p<0.009, ****p<0.0001, 16-22 mice per genotype). **E**. Comparison of survival time after KA administration between WT and R/+ mice (****p<0.0001, 16-22 mice per genotype). **F**. Heat maps depicting individual level seizure severity after KA administration organized by genotype. Individual mice are presented on the y-axis, and the x-axis represents 5 min time bins.

*Atp1a3*^*R/+*^ mice exhibited lower seizure thresholds evoked by two additional induction methods. Immediately following a subthreshold electroconvulsive shock (ECS; 38 mA), few (2/10) WT mice but most (8/9) *Atp1a3*^*R/+*^ mice exhibited tonic hindlimb extension (HLE; p<0.0001, Fig. 5C). In response to kainic acid (KA) exposure, *Atp1a3*^*R/+*^ mice progressed faster to Racine stage 3 (p<0.009) and stage 4 (p<0.0001) with an average latency of 17.9 ± 2.3 and 25.2 ± 3.0 min, respectively, compared to 30.1 ± 2.8 and 51.6 ± 15.3 min observed for WT mice (Fig. 5D). Furthermore, none of the *Atp1a3*^*R/+*^ mice (0/16) survived post KA exposure (median survival 29.9 min), while 81.8% (18/22) of WT mice survived (p<0.0001; Fig. 5E).

We video monitored *Atp1a3*^*R/+*^ mice for 1 hour to detect handling-induced seizure activity. We did not observe seizures in 13 WT mice, but 38% (5/13) of *Atp1a3*^*R/+*^ mice exhibited post-handling seizures (p<0.04; Supplemental Fig. S3). Of *Atp1a3*^*R/+*^ mice with seizures, 3 had a single GTCS with varying degrees of severity, including one that ended in HLE followed immediately by death, which resembles sudden unexplained death in epilepsy (SUDEP) observed in some mouse models of developmental and epileptic encephalopathy (24, 25).

### Atp1a3^G947R^ mice exhibit spontaneous seizures and abnormal EEG activity

To further investigate whether *Atp1a3*^*R/+*^ mice had unprovoked seizures and epilepsy, we performed continuous video-EEG recording over multiple days to quantify spontaneous seizures in WT and *Atp1a3*^*R/+*^ mice. Blinded manual review of the data revealed significant differences in EEG activity between genotypes. Spontaneous seizures were detected by extended EEG recordings in two *Atp1a3*^*R/+*^ mice (Table 3, Fig. 6A). Corresponding to the electrographic activity, *Atp1a3*^*R/+*^ mice exhibited behavioral arrest and/or head bobbing and then returned to normal activity after seizure termination. In addition to spontaneous seizures and EEG power alterations, multiple instances of high amplitude, high frequency bursting and low frequency (< 3 Hz) spike train activity were detected in *Atp1a3*^*R/+*^ mice (Fig. 6B,C). *Atp1a3*^*R/+*^ mice exhibited recurrent high-amplitude, high-frequency spike bursts significantly more frequently than WT mice (Table 3). No difference in baseline activity, calculated by the root mean square (RMS) was detected by genotype (Table 3; Fig. 6D). In approximately 500 hours of analyzed EEG traces, 40% (2/5) of WT mice exhibited rare bursts, with a median daily frequency of 0 segments per day (95% CI [0-0.8]; Table 3). In contrast, 100% (11/11) of *Atp1a3*^*R/+*^ mice exhibited bursts, with a median daily frequency of 4.3 segments per day (95% CI [0.3-35.2]) (p<0.02; Fig. 6E). Similarly, spike trains were observed in 91% (10/11) of *Atp1a3*^*R/+*^ mice during approximately 1100 hours of analyzed EEG traces, with a median duration of 347 seconds (95% CI [261-448]; Table 3). No spike trains were observed in WT mice during 500 hours of analyzed EEG traces (Fig. 6F, Supplemental Fig. S4).

**Table 3.** EEG Characteristics of individual mice.

| Mouse | | Manually<br>Reviewed<br>Hours | RMS<br>(μV) | Daily<br>Burst<br>Frequency | Number of<br>Trains<br>(duration)<sup>a</sup> | Features |
| --- | --- | --- | --- | --- | --- | --- |
| WT | 1 | 167.2 | 16.94 | 0.0 | 0 (0) | unremarkable |
|  | 2 | 96.5 | 36.07 | 0.7 | 0 (0) | unremarkable, bursts |
|  | 3 | 75.7 | 22.72 | 0.3 | 0 (0) | unremarkable, bursts |
|  | 4 | 75.6 | 24.7 | 0.0 | 0 (0) | unremarkable |
|  | 5 | 82.9 | 27.89 | 0.0 | 0 (0) | unremarkable |
| R/+ | 1 | 96.7 | 9.18 | 35.2 | 23 (335) | trains, bursts |
|  | 2 | 72.8 | 19.45 | 4.3 | 0 (0) | bursts |
|  | 3 | 73.0 | 48.69 | 0.7 | 6 (38) | trains, bursts |
|  | 4 | 75.5 | 50.62 | 3.5 | 5 (93) | trains, bursts |
|  | 5 | 72.9 | 18.65 | 5.9 | 0 (0) | seizures, bursts |
|  | 6 | 73.1 | 24.53 | 6.9 | 0 (0) | bursts |
|  | 7 | 96.5 | 51.41 | 0.7 | 0 (0) | unremarkable, bursts |
|  | 8 | 239.6 | 18.44 | 41.2 | 20 (123) | seizures, trains, bursts |
|  | 9 | 96.6 | 24.45 | 6.2 | 1 (1.4) | trains, bursts |
|  | 10 | 95.9 | 24.63 | 0.3 | 20 (154) | trains, bursts |
|  | 11 | 96.0 | 16.28 | 13.5 | 6 (73) | trains, bursts |

<sup>a</sup> total duration in minutes given in parentheses

**Fig. 6.**
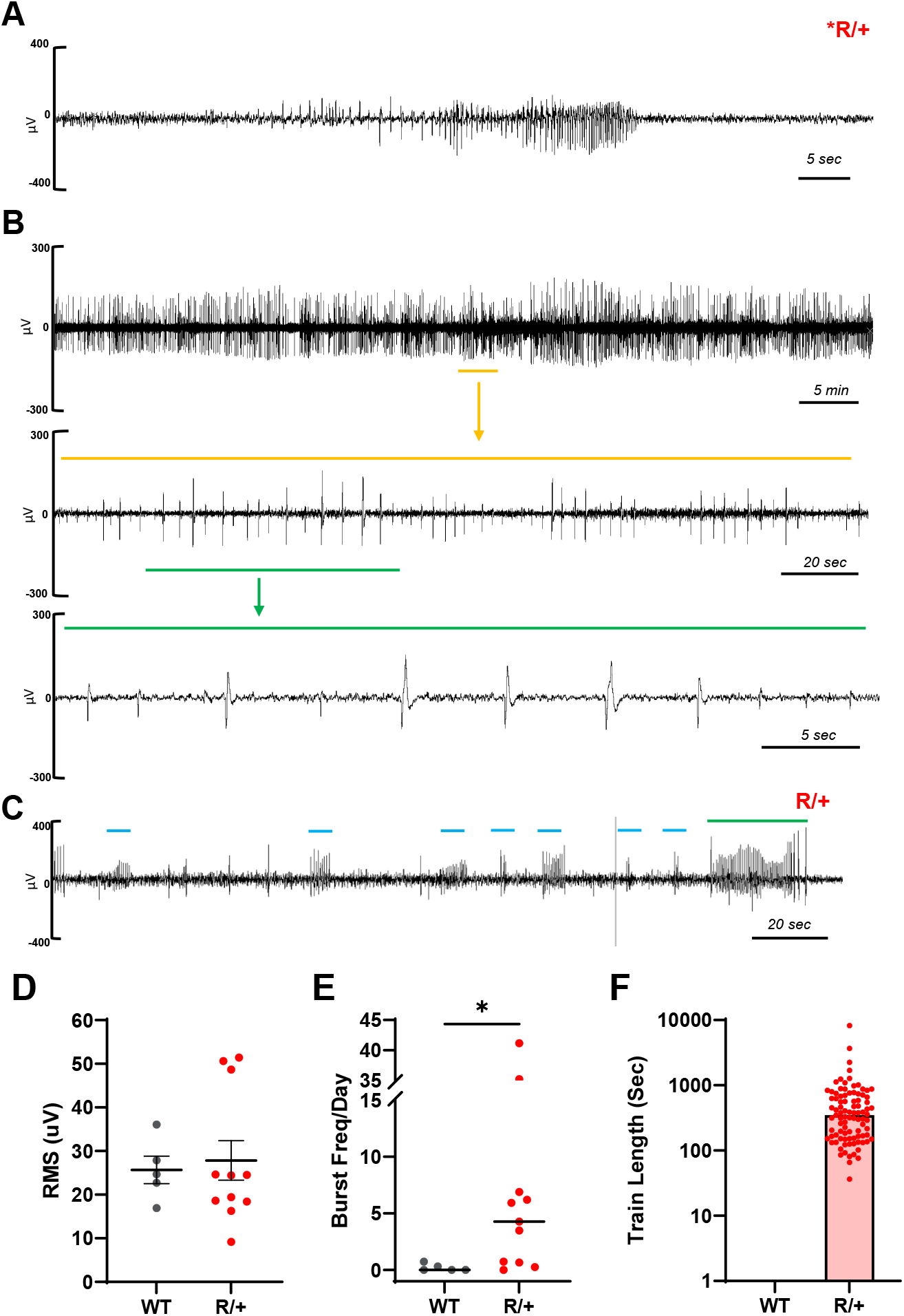
Heterozygous *Atp1a3*^G947R^ mice have spontaneous epileptiform EEG activity. **A**. Representative spontaneous seizure recorded by EEG from a R/+ mouse. Corresponding to the electrographic activity, this R/+ mouse exhibited behavioral arrest and returned to normal activity after seizure termination. **B**. Segment (∼1 h) of a continuous spike train identified in a R/+ mouse. Yellow line represents a ∼3.5 min segment expanded into middle trace. Green line represents a ∼40 s segment expanded into bottom trace. **C**. Representative trace of a R/+ mouse with frequent high amplitude, high frequency bursts (blue lines) and spike train (green line). **D**. Comparison of root square mean (RMS) between WT and R/+ mice (p>0.76, 5-11 mice per genotype). **E**. Comparison of burst frequency between WT and R/+ mice (*p<0.02, 5-11 mice per genotype). **F**. Duration of spike trains in WT and R/+ mice (5-11 mice per genotype).

We used power spectral density (PSD) analysis to quantify the power of different frequency components to discern the functional state of the *Atp1a3*^*R/+*^ brain during periods of apparent sleep (no EMG activity) and wakefulness (Fig. 7). *Atp1a3*^*R/+*^ mice exhibited significantly lower spectral power during apparent sleep, particularly in the 3.1-9.4 Hz frequency range (upper delta, theta and lower alpha bands), compared to WT mice (p<0.03-0.0001; Fig. 7A). Furthermore, the average total power during apparent sleep was significantly lower in *Atp1a3*^*R/+*^ mice, with an average of 389.9 ± 51.3 μV^2^ compared to the WT average of 816.2 ± 105.6 μV^2^ (p<0.007; Fig. 7B). During wakefulness, *Atp1a3*^*R/+*^ mice also showed lower spectral power within the 4.7-9.4 Hz frequency range (theta and lower alpha bands) compared to WT mice (p<0.002-0.0001; Fig. 7C). Furthermore, the total power during wakefulness was significantly lower in *Atp1a3*^*R/+*^ mice, 186.3 ± 24.6 μV^2^ compared to WT mice, 347.5 ± 47.3 μV^2^ (p<0.02; Fig. 7D).

**Fig. 7.**
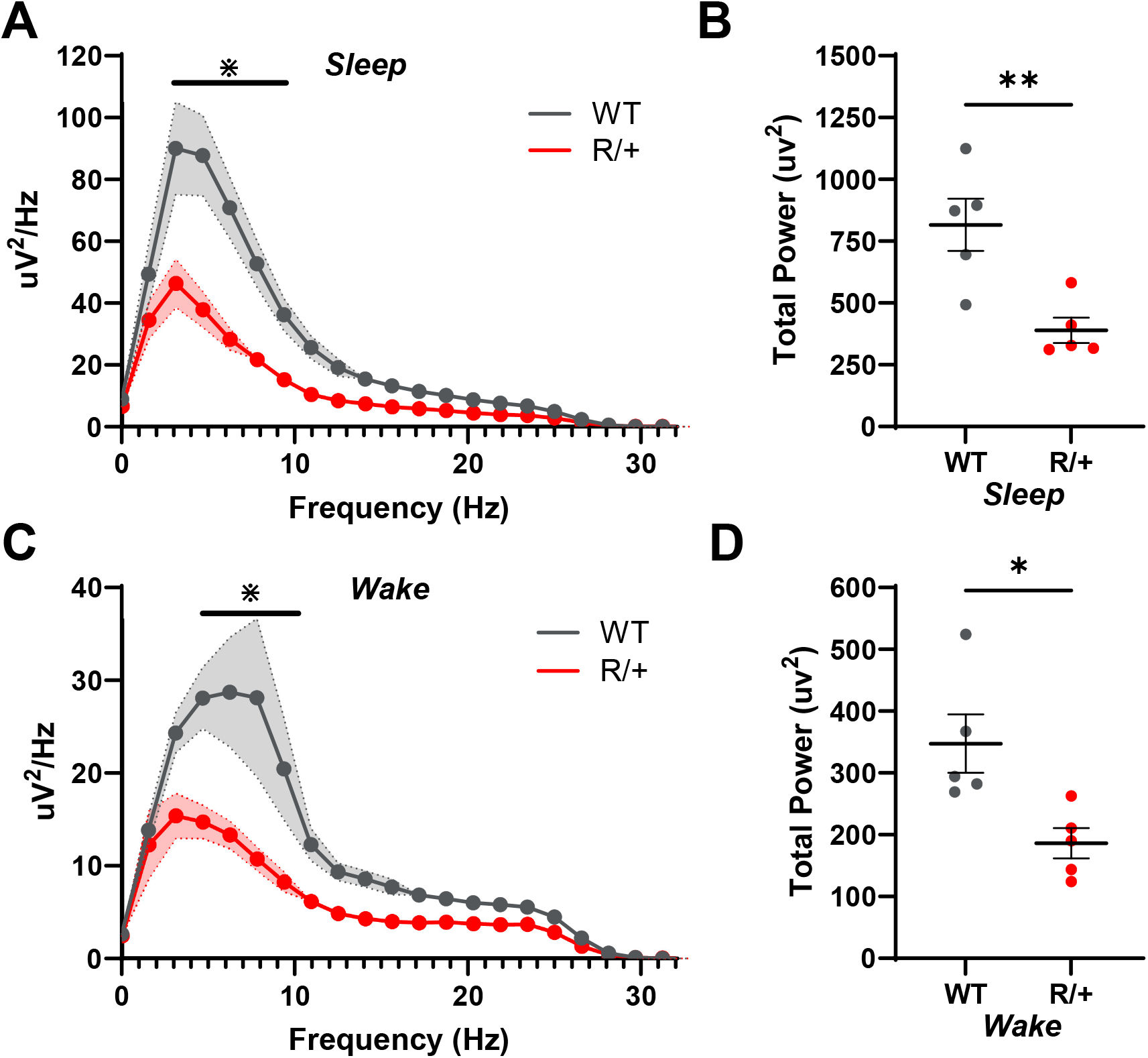
Power spectral density (PSD) analysis. **A**. R/+ mice exhibit lower activity across the power spectra during apparent sleep (3.1-9.4 Hz frequency range: *p<0.03-0.0001). Data points represent mean values from 5 mice per genotype and shading indicate SEM. **B**. Comparison of total power between WT and R/+ mice during apparent sleep (**p<0.007, 5 mice per genotype). **C**. R/+ mice exhibit lower activity across the power spectra during wakefulness (4.7-9.4 Hz frequency range: *p<0.002-0.0001). Data points represent mean values from 5 mice per genotype and shading indicate SEM. **D**. Comparison of total power between WT and R/+ mice during wakefulness (*p<0.02, 5 mice per genotype). PSDs and total power were averaged from artifact, spike, and seizure free 5-minute epochs. In panels B and D, symbols are data from individual mice, horizontal lines represent mean values and error bars indicate SEM.

## DISCUSSION

We generated and investigated a new *Atp1a3*^G947R^ mouse model of the rare, neurodevelopmental disorder AHC caused by a recurrent pathogenic missense variant in *ATP1A3*. Observed phenotypic features of this novel AHC model compared with WT mice included small body size, lower long-term survival, hyperactivity, anxiety-like behaviors, severe motor dysfunction with low grip strength, impaired coordination with abnormal gait and balance, cooling-induced hemiplegia and dystonia, lower seizure threshold, spontaneous seizures, sudden death after seizures, and abnormal EEG activity. These observations resemble core clinical features of human AHC, which establishes face validity of the *Atp1a3*^G947R^ mouse model.

Many of the features observed in heterozygous *Atp1a3*^G947R^ mice are shared with two previously reported mouse models of AHC with other recurrent human *ATP1A3* mutations (D801N, E815K). The availability of all three models represents the spectrum of disease severity more fully and provides new opportunities to identify shared pathophysiological mechanisms. An important distinction between the previously reported AHC mouse models and our study relates to genetic background. The two reported AHC models representing other recurrent mutations and the *Myk* mouse were generated using homologous recombination or by chemical mutagenesis on mixed strains. Both D801N and E815K mice were created using 129Sv embryonic stem cells then crossed to C57BL/6J (15, 16). Our *Atp1a3*^G947R^ is the first mouse model of AHC created using gene editing and the first generated on a pure genetic background.

In addition to the neurological features, we also observed non-Mendelian offspring genotype ratios during conventional breeding of founder animals on the C57BL/6J genetic background. However, IVF and breeding to a different strain were successful in generating offspring with near-Mendelian ratios suggesting that poor breeding capability was not related to embryonic lethality of the mutation. We speculate that this problem could be related to impaired sperm function or reproductive fitness of *Atp1a3*^G947R^ C57BL/6J males. Another murine Na/K-ATPase (*Atp1a4*) is essential for sperm motility (26) but how this relates to *Atp1a3*^G947R^ is unclear. We intentionally screened for potential off-target genome edits in *Atp1a4* to exclude a molecular disruption of this gene. Interestingly, prior to the use of CRISPR/Cas9 gene editing to generate mutant mice, embryonic stem (ES) cells from C57BL/6J were seldom utilized in gene targeting due to the known technical and biological challenges of the strain (27).

Seizures occur in a large proportion of individuals affected with AHC. In large case series of AHC, 40 -50% were reported to have co-morbid epilepsy (3, 4). Indeed, seizures may be among the earliest paroxysmal feature of the disease, which can be severe and refractory to treatment (6, 28). Among the most recurrent AHC-associated *ATP1A3* mutations, seizures in individuals harboring the E815K mutation are most severe. Seizure frequency is lower in individuals with the G947R mutation, but onset of epilepsy may be earlier (4). Given the high frequency and treatment-refractory nature of epilepsy in AHC, well-characterized *in vivo* model systems that recapitulate this disease feature in a quantitative manner will have value for testing new therapeutic strategies.

Three previously reported mouse models of AHC exhibited seizures especially upon handling (13, 15, 16). Epilepsy in the *Myk* mouse is strain-dependent. On the mixed (129S1/SvImJ | C57BL/6N) background, *Myk* mice exhibited a prominent and severe epilepsy phenotype that ultimately disappeared when mice were bred > 20 generations to C57BL/6NCr (12, 29). The identities of the presumed genetic modifiers responsible for the strain differences of the *Myk* epilepsy phenotype are unknown. Heterozygous D801N mice exhibit enhanced seizure susceptibility to electrical kindling of the amygdala and with flurothyl exposure, and also exhibited spontaneous and/or handling-induced behavioral seizures (15, 30). Similarly, mice heterozygous for *Atp1a3*-E815K exhibited spontaneous and induced seizures with associated high mortality (16). EEG analyses were limited for these previously reported AHC mouse models.

Our study of heterozygous *Atp1a3*^G947R^ mice provides an in-depth electrographic analysis of seizures in an AHC mouse model. In addition to scoring spontaneous and induced seizures, we quantified EEG spectral power during apparent sleep and wake states. We observed significantly reduced total EEG power in the mice, which is a marker of diminished brain activity and connectivity (31). These EEG findings are quantifiable electrophysiological biomarkers of epilepsy and could be used as readouts for evaluating preclinical efficacy of antiseizure therapies.

## CONCLUSIONS

In summary, our study demonstrated the neurological phenotypes exhibited by a novel mutant mouse model of the rare neurodevelopmental disorder AHC associated with the recurrent *ATP1A3* pathogenic variant p.G947R. Our new mouse model reproduces cardinal features of human AHC including inducible dystonia and hemiplegia along with a prominent seizure phenotype, including lower thresholds for chemically and electrically induced seizures, and spontaneous behavioral and electrographic seizures. Our quantitative EEG analysis provides novel insights about comorbid epilepsy in AHC that were not apparent in previously reported mouse models of this disorder. This new mouse model will have value as a platform for investigating AHC pathophysiology and for *in vivo* testing of new therapeutic strategies.

## ACKNOWLEDGEMENTS

We thank Levi Barse, Tyler Thenstedt, Rylie Pancoast, Kelsey Davis and Olivia Valente for technical assistance. The genetically engineered mice were generated with the assistance of Northwestern University Transgenic and Targeted Mutagenesis Laboratory.

## Supplemental Information

### Supplemental Tables

**Table S1.**
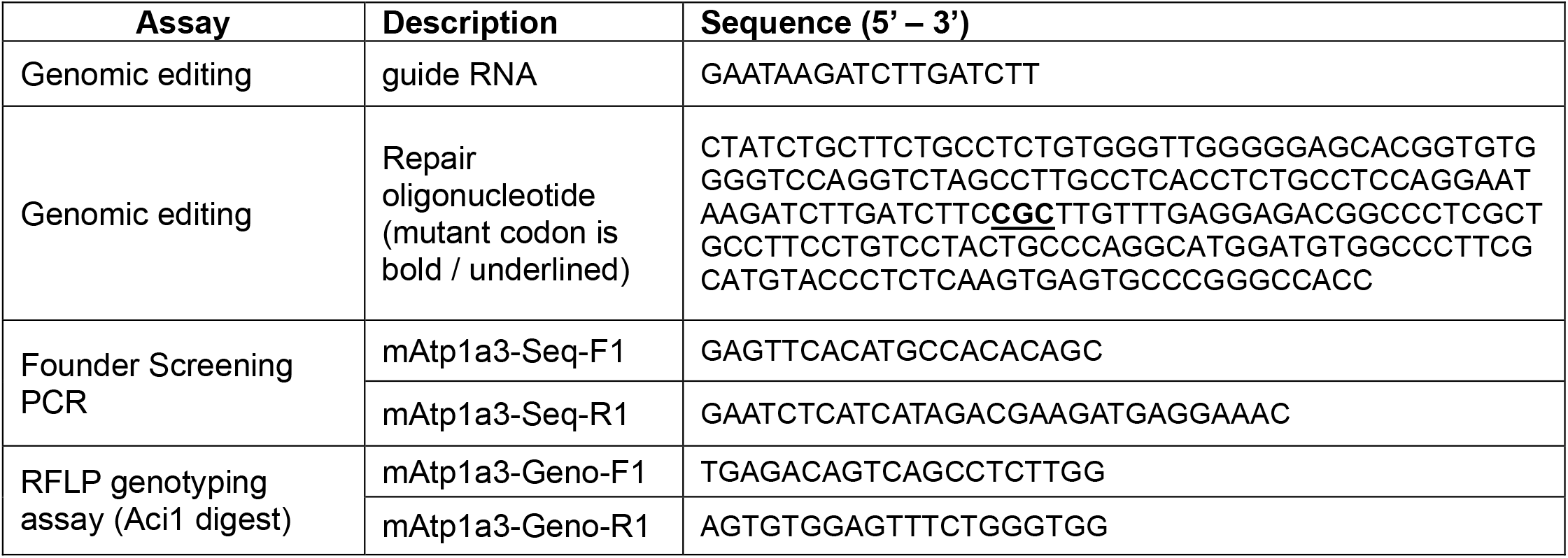
Primer sequences used for genome editing and genotyping.

**Table S2.** Experimental cohorts.

| Cohort | IVF | Experiments | Age (P) |
| --- | --- | --- | --- |
| 1 | 2021 | Seizure monitoring | 37-46 |
|  |  | Stride and rotarod | 48-52 |
|  |  | Expression | 52-55 |
| 2 | 2022 | Open field | 88-101 |
|  |  | Tail suspension | 101-113 |
|  |  | Forelimb swipe | 101-113 |
|  |  | Wire hang | 101-113 |
|  |  | Balance beam | 107-116 |
|  |  | Body weight/length | 107-116 |
|  |  | EEG | 113-151 |
| 3 | 2023 | Body weight/length | 79-86 |
| 4 | 2024 | Dystonia | 98-108 |
|  |  | ECS | 103-109 |
| 5 | 2024 | Flurothyl | 90-119 |
| 6 | 2024 | KA | 97-108 |
|  |  | Weight | 97-108 |
| 7 | 2024 | EEG | 100-110 |
|  |  | Body weight | 100-110 |

**Table S3.** Statistical methods.

| Figure | Comparison | Test | Value | Post Hoc |
| --- | --- | --- | --- | --- |
| 1B | Transcript | Unpaired T | p>0.43 | n/a |
| 1C | Protein | Mann-Whitney | p=0.31 | n/a |
| 2A | Weight (by sex) | Unpaired T | p<0.0001 | n/a |
| 2B | Length (by sex) | Unpaired T | p<0.0001 | n/a |
| 2C | Survival | Mantel-Cox | P<0.0001 | n/a |
| 3A | Open field distance | Unpaired T | P<0.0001 | n/a |
| 3B | Open field % center time | Unpaired T | p<0.03 | n/a |
| 3C | Tail suspension | Mann-Whitney | p<0.0001 | n/a |
| 3D | Forelimb swipe | Mann-Whitney | p<0.0001 | n/a |
| 3E | Balance beam | Fishers Exact | p<0.03 | n/a |
| 3F | Wire hang | Mantel-Cox | p<0.0002 | n/a |
| 4A | Cooling survival | Mantel-Cox | p<0.004 | n/a |
| 4B | Recovery | Mann-Whitney | p<0.0001 | n/a |
| 4C | Dystonia | Mann-Whitney | p<0.0001 | n/a |
| 5A | Flurothyl GTCS | Mann-Whitney | p=0.0001 | n/a |
| 5B | Flurothyl interval | Mann-Whitney | p<0.0001 | n/a |
| 5C | ECS | Fishers Exact | p<0.0001 | n/a |
| 5D | KA Stage | Two-way ANOVA | F(1,129)=23.3;<br>p<0.0001 | Sidak's;<br>p<0.009,p<0.0001 |
| 5E | KA survival | Mantel-Cox | P<0.0001 | n/a |
| 6A | PSD *sleep*: 3.1 Hz* | Two-way ANOVA | F(1,176)=84.3;<br>p<0.0001 | Sidak's; p<0.0001 |
|  | PSD *sleep*: 4.7 Hz* |  |  | Sidak's; p<0.0001 |
|  | PSD *sleep*: 6.3 Hz* |  |  | Sidak's; p<0.0001 |
|  | PSD *sleep*: 7.8 Hz* |  |  | Sidak's; p<0.0001 |
|  | PSD *sleep*: 9.4 Hz* |  |  | Sidak's; p<0.03 |
| 6B | Total power *sleep* * | Unpaired T | p<0.007 | n/a |
| 6C | PSD *wake*: 4.7 Hz | Two-way ANOVA | F(1,176)=54.3;<br>p<0.0001 | Sidak's; p<0.0003 |
|  | PSD *wake*: 6.3 Hz |  |  | Sidak's; p<0.0001 |
|  | PSD *wake*: 7.8 Hz |  |  | Sidak's; p<0.0001 |
|  | PSD *wake*: 9.4 Hz |  |  | Sidak's; p<0.002 |
| 6D | Total power *wake* | Unpaired T | p<0.02 | n/a |
| 7D | RMS | Unpaired T | p>0.76 | n/a |
| 7E | Burst frequency | Mann-Whitney | p<0.02 | n/a |

\*apparent sleep period

### Supplemental Figures

**Fig. S1.**
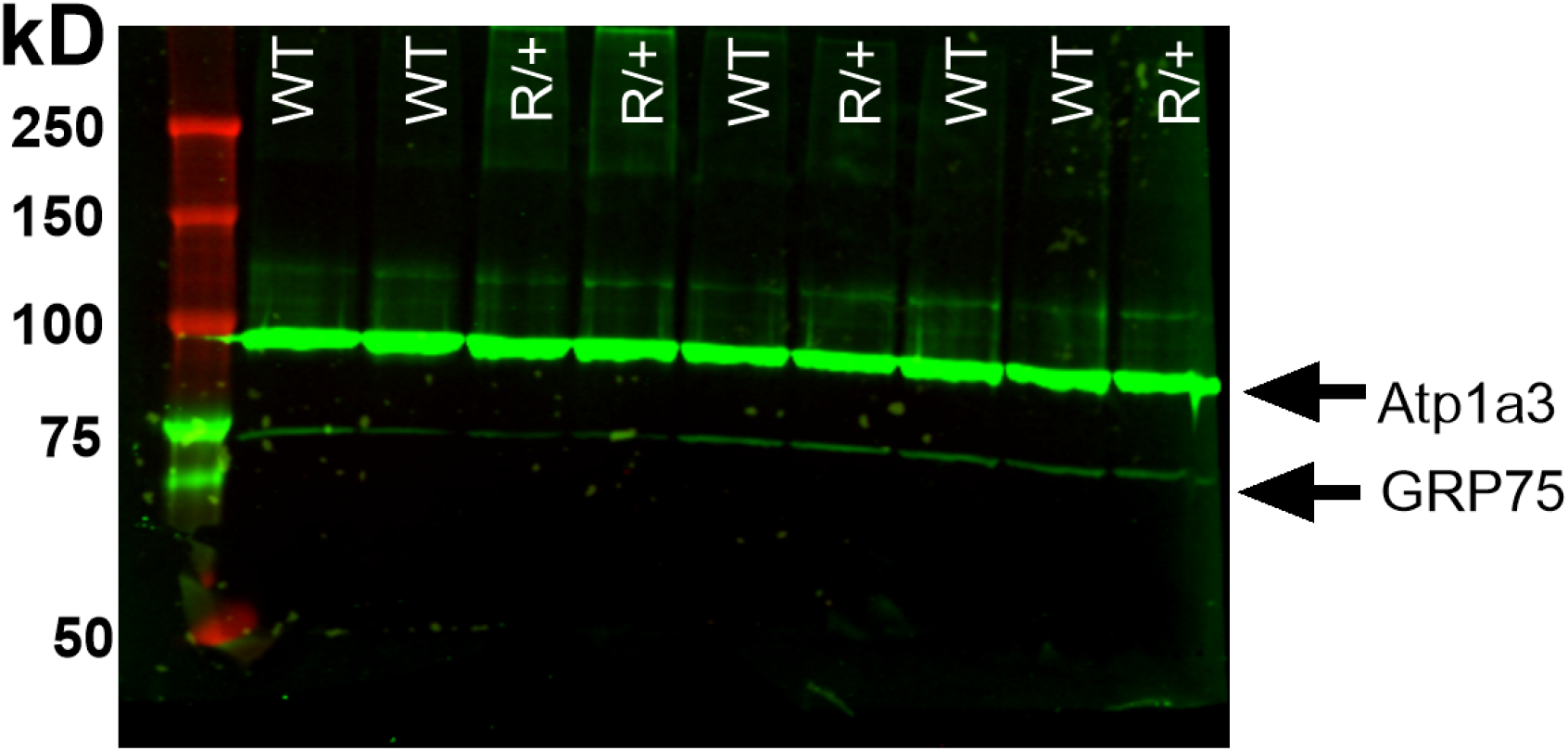
Western blot analysis of WT and heterozygous Atp1a3-G947R mice. Representative western blot demonstrating no difference in protein expression between genotypes. Each lane represents protein from whole brain lysates from individual WT or R/+ mice. Locations of Atp1a3 and GRP75 proteins are indicated by arrows. Molecular mass standards are in the first lane, labeled by kD on the left.

**Fig. S2.**
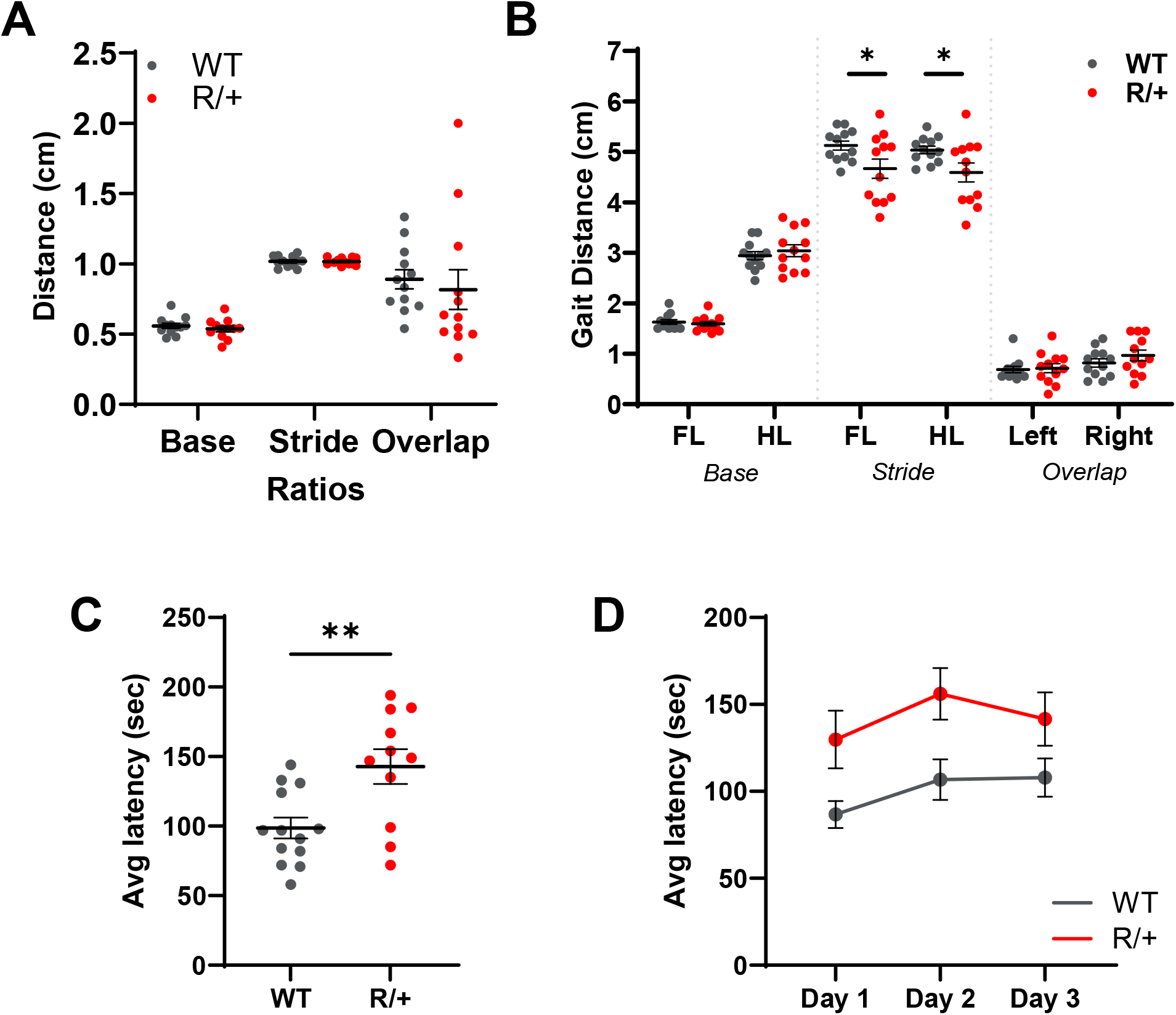
Gait and rotarod performance. **A**. Analysis of gait ratios demonstrated no differences in base, stride or overlap were detected between WT and R/+ mice (one-way ANOVA F(1,66)=0.36, p>0.54, 12 mice per genotype). **B**. R/+ mice have shorter forelimb (FL) and hindlimb (HL) stride distances compared to WT, while no difference in base or overlap gait were detected. WT had an average FL and HL stride distance of 5.1 ± 0.1 and 5.0 ± 0.07 cm, while R/+ had an average FL and HL stride distance of 4.7 ± 0.2 and 4.6 ± 0.2 cm (FL: *p<0.02, HL: *p<0.03, 12 mice per genotype). **C**. Average latency to fall from accelerating rotarod. Mutant mice exhibited a longer fall latency of 142.8 ± 12.5 sec compared to the WT average of 98.6 ± 7.4 sec (unpaired t-test, **p<0.005, 11-13 mice per genotype). **D**. Daily trial data for average fall latencies from accelerating rotarod. Through all trial days, R/+ mice consistently remained on the accelerating rotarod for longer durations compared to WT. *Day 1* WT: 86.7 ± 7.8; R/+: 129.8 ± 16.6. *Day 2* WT: 106.8 ± 11.6; R/+: 156.1 ± 14.8. *Day 3* WT: 107.9 ± 11.0; R/+: 141.5 ± 15.3 (11-13 mice per genotype).

**Fig. S3.**
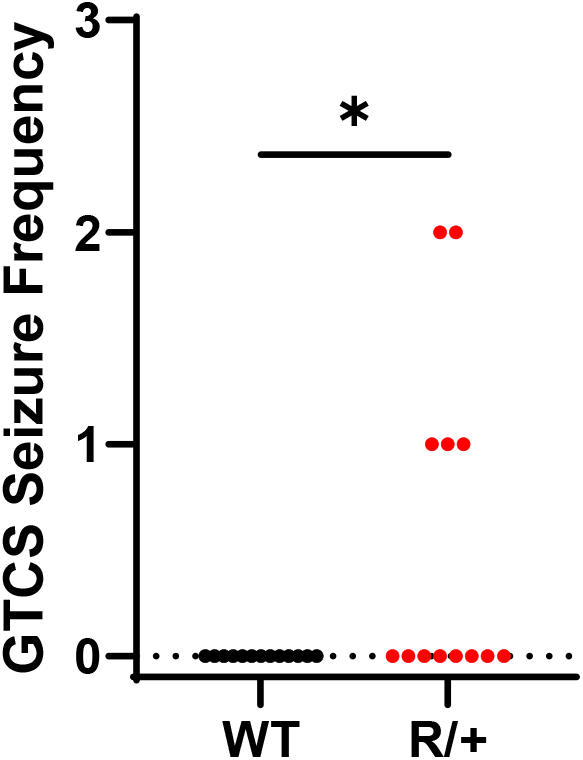
Post-handling seizure frequency. WT and R/+ mice were video-monitored for one hour post handling. GTCS activity was identified in 38% (5/13) of R/+ mice post handling. WT mice exhibited no observable seizure activity. (*p<0.02, 13 mice per genotype). Symbols are data from individual mice.

**Fig. S4.**
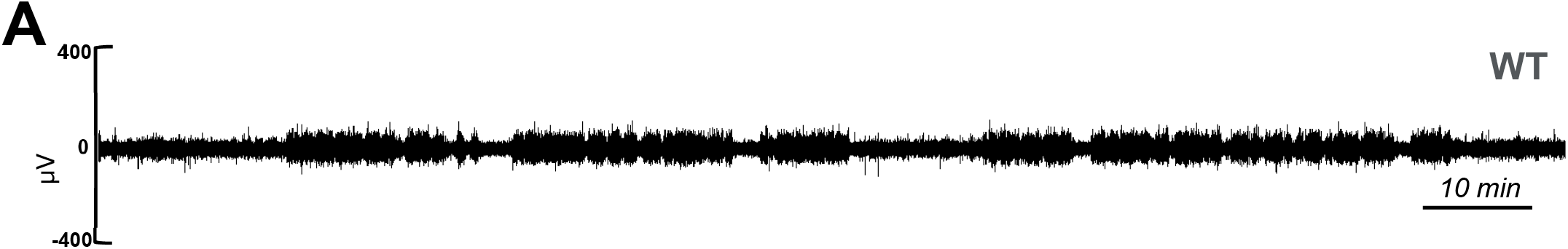
Extended EEG recording from a WT mouse. Representative ∼2 h EEG trace from a WT mouse exhibited unremarkable features.

